# Lineage tracing of Notch1-expressing cells in intestinal tumours reveals a distinct population of cancer stem cells

**DOI:** 10.1101/364349

**Authors:** Larissa Mourao, Guillaume Jacquemin, Mathilde Huyghe, Wojciech J. Nawrocki, Naoual Menssouri, Nicolas Servant, Silvia Fre

## Abstract

Colon tumours are hierarchically organized and contain multipotent self-renewing cells, called Cancer Stem Cells (CSCs). We have previously shown that the Notch1 receptor is expressed in Intestinal Stem Cells (ISCs); given the critical role played by Notch signalling in promoting intestinal tumourigenesis, we explored Notch1 expression in tumours. Combining lineage tracing in two tumour models with transcriptomic analyses, we found that Notch1 + tumour cells are undifferentiated, proliferative and capable of indefinite self-renewal and of generating a heterogeneous clonal progeny. Molecularly, the transcriptional signature of Notch1+ tumour cells highly correlates with ISCs, suggestive of their origin from normal crypt cells. Surprisingly, Notch1+ expression labels a subset of CSCs that show reduced levels of Lgr5, a reported CSCs marker. The existence of distinct stem cell populations within intestinal tumours highlights the necessity of better understanding their hierarchy and behaviour, to identify the correct cellular targets for therapy.

## Introduction

Intestinal crypts have been reported to harbour two distinct types of stem cells: homeostatic stem cells, marked by the G-protein coupled receptor Lgr5^1^, that continuously generate new progenitors to ensure efficient renewal of the intestinal mucosa, and presumably quiescent stem cells, thought to provide a reserve source of stem cells that can be activated upon injury^2,3^. We have shown that the Notch1 receptor is expressed in both homeostatic and reserve stem cells populations *in vivo*, providing a valuable tool to mark both cell types and dissect their hierarchical relationship^4^. Intestinal tumours have been proposed to originate from intestinal stem cells (ISCs) or very early progenitors, as these are the only cells that persist long enough within the tissue to ensure clonal expansion of a mutant progeny^5^. However, it should be noted that differentiated cells and/or intestinal progenitors have been shown to be able to undergo dedifferentiation upon activation of specific pathways and to acquire stem cell properties that may result in tumour formation^6^. However, these studies are based on genetic targeting of specific oncogenic mutations, while direct evidence showing the presence of CSCs within spontaneously arising intestinal tumours is still poor. Recently, a lineage tracing study in intestinal and colonic tumours using two stem cell-specific promoters, Lgr5 and Bmi1^7^, proposed that these two ISCs markers can be used to detect different stem cell populations in intestinal tumours^8^. We have previously shown that the specific expression of the Notch1 receptor in ISCs promotes active Notch signalling in these cells^4^ and, consistently with its essential role in intestinal homeostasis and cell fate determination^9^, Notch activity has been reported as essential for crypt stem cells maintenance^10^. Of relevance, Notch signalling is required for intestinal tumour formation and it cooperates with the Wnt pathway to promote crypt hyperplasia^11,12^. On these premises, we assessed whether the expression of the Notch1 receptor labels CSCs in intestinal and colon adenomas derived from both a genetic mouse model and a carcinogenic protocol. We performed clonal analysis using the Notch1-Cre^ERT2^ mouse line, which we have previously shown to mark multipotent ISCs in the small intestine and colon^4^, and assessed the identity and fate of Notch1+ cells within spontaneously arising intestinal tumours.

We show that Notch1-labeled cells represent a distinct population of CSCs within both intestinal and colon tumours, which contributes to tumour growth by clonal expansion and generates intra-tumoural heterogeneity. These *in vivo* studies provide evidence for the existence of different types of CSCs in intestinal tumours, which might have different origins and/or exhibit differential response to treatment.

## Results

### Notch1Cre^ERT2^ labels undifferentiated and proliferative tumour cells

To track Notch1+ intestinal adenoma cells *in vivo*, we generated a triple transgenic mouse (hereafter referred to as N1-Cre/mTmG/Apc) by crossing mice carrying both the Notch1-Cre^ERT2^ (N1-Cre)^4^ and the Rosa26^mTmG^ (mTmG)^13^ reporter with Apc+^/1638N^ (Apc) mutant mice^14^ (Fig. 1a). Apc mice harbour a heterozygous germline mutation in the *Apc* tumour suppressor gene and spontaneously develop intestinal adenomas, initially detectable at around six months of age, due to loss of heterozygosity (LOH) at the *Apc* locus. In our compound N1-Cre/mTmG/Apc mice, the membrane-associated red fluorescent protein (mT) is expressed in all cells, while membrane-associated GFP (mG) marks Cre-targeted cells. To identify the cells expressing the Notch1 receptor within tumours, N1-Cre/mTmG/Apc tumour-bearing mice received a single dose of tamoxifen and were analysed 24h later (Fig. 1b). Quantification by flow cytometry of the proportion of Notch1+ cells within tumour epithelial cells (referred as TEC, selected with the markers EpCAM+/Lin-, gate strategies are detailed in Supplementary Fig. 1), indicated, in agreement with our immunofluorescence results, that Notch1-expressing epithelial cells represent a rare tumour cell population comprising 1,2% ± 0,3% of TEC (Fig. 1c). It should be noted that, as the N1-Cre line also labels other types of stromal cells, we exclusively focused our analysis on epithelial cells, expressing the epithelial marker EpCAM (Epithelial cell adhesion molecule^15^ (Fig. 1d). Since Apc mutant intestinal tumours present differentiated tumour cells, we evaluated if Notch1 is expressed in such cells by immunostaining for differentiation markers for secretory cells, such as Agglutinin (Ulex Europaeus Agglutinin, labelling both Paneth and Goblet cells), Lysozyme1 (a specific marker of Paneth cells)^16^ and Mucin2 (expressed in Goblet cells^17^) (Fig. 1d). None of these markers was expressed in GFP+ cells, consistently with the lack of Notch1 expression in secretory cells in the normal intestinal epithelium^9^. We also assessed the expression of secretory and enterocyte (Alkaline Phosphatase Intestinal, Alpi^18^) markers by qRT-PCR on sorted tumour cells and confirmed that GFP+ cells show strongly reduced levels of expression for all of these markers (Fig. 1e), indicating that the N1-Cre mouse line labels undifferentiated tumour cells.

**Figure 1.**
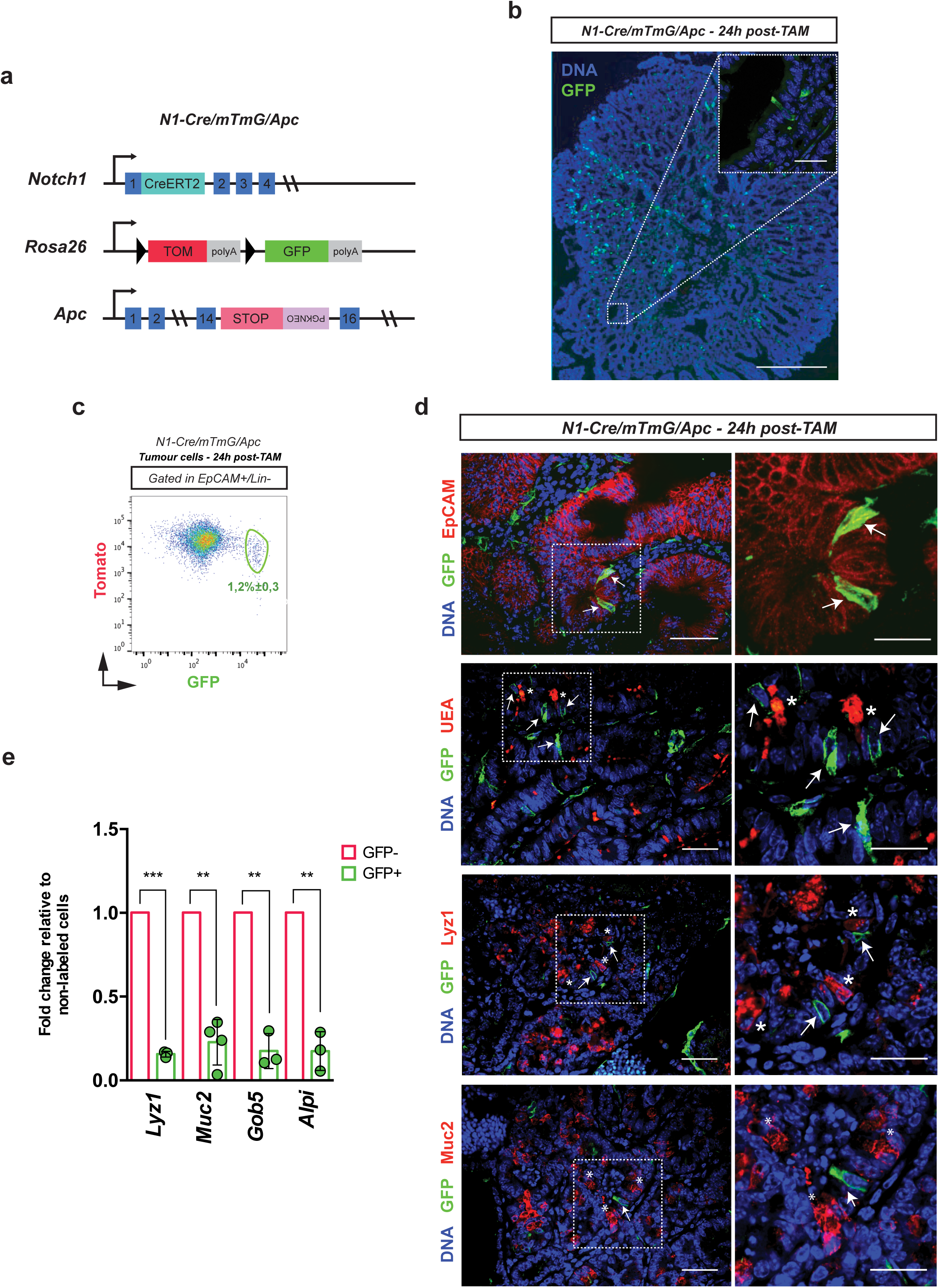
Notch1Cre^ERT2^ labels undifferentiated and proliferative tumour cells. **(A)** Schematic representation of the triple transgenic mouse model used in this study. Notch1-CreERT^SAT^ knock-in mice (referred to as N1-Cre) were crossed to Rosa26mT/mG reporter mice (named mTmG) and to Apc^+/1638N^ mice (termed Apc). **(B)** Representative section of an intestinal tumour from N1-Cre/mTmG/Apc mice, 24h post tamoxifen injection. The inset shows a higher magnification of a Notchl-expressing tumour cell (marked by GFP in green). DNA is labelled by DAPI in blue. Scale bars represent 200μm and 15μm in the magnification panel. **(C)** FACS analysis (see Supplementary Fig. 1a for gate strategy details) of N1-Cre/mTmG/Apc dissociated tumour cells 24h post induction. Lin+ cells were excluded and single epithelial tumour cells were gated as TEC (Epcam+/Lin-), allowing the quantification of Notch1+ tumour cells. Note that GFP+ cells also display Tomato fluorescence 24h after induction (GFP+/Tom+), as the Tomato protein is still present at this time point, even if recombination has occurred. **(D)** Immunofluorescence stainings of N1-Cre/mTmG/Apc tumour sections using anti-EpCAM, Agglutinin (UEA), anti-lysozyme (Lyz1) and anti-Mucin2, all in red. Notch1-expressing tumour cells are labelled in green (GFP+) and DNA is marked by DAPI in blue. Magnifications insets are shown in the right panels. Arrows indicate Notch1-expressing tumour cells and asterisks show secretory tumour cells. **(E)** qRT-PCR showing the relative RNA expression of Lyz1, Muc2, Gob5 and Alpi in Notch1+ (green bars) and non-recombined tumour cells (red bars). Bars represent the average ± SDs of independent biological replicates (n ≥ 3) normalized to the 18S housekeeping gene. **= P ≤ 0.005; *= P ≤ 0.05. The p-values were calculated using the Paired Ratio t-test. Scale bars correspond to 15μm in the inset in **(B)**, 30μm in **(D)** and 20μm in the insets in **(D)**.

### Notch1 expression defines multipotent tumour cells with self-renewal capacity

To map the fate of Notch1+ tumour cells and establish their self-renewal capacity *in vivo*, we examined tumour-bearing N1-Cre/mTmG/Apc mice at different time points after administration of tamoxifen, from 4 days up to 90 days (Fig. 2a). Our experimental approach required that we started tracing in tumour-bearing mice (thus in 6-month old animals, which generally succumb 2 to 3 months later) and consequently, a 90 days chase was the longest chase period for clonal analysis. Lineage tracing showed that Notch1+ tumour cells rapidly generate clones of marked progeny, visible already 4 days after labelling, and that these clones enlarge over time (Fig. 2b). To quantify the clonal expansion of Notch1 lineages *in vivo* in growing tumours, we used two different methods: by FACS analysis, we assessed the percentage of GFP+ cells within total TEC at each time point (Supplementary Fig. 2a and 2b); by immunofluorescence, we quantified the GFP+ tumour area within the total tumour surface, defined by the expression of the Tomato fluorescent protein (Supplementary Fig. 2c). From 24h post-tamoxifen up to 90 days, the proportion of GFP+ tumour cells quantified by FACS increased from 1,2% ± 0,3% to 6,1% ± 1,95% within total TEC (Fig. 2c). A two-parameter logarithmic function yielded the best fit to describe both quantification methods. This function predicts that Notch1+ lineages will persist in adenomas and continue to slowly expand even after 90 days, indicating self-renewal capacity (Fig. 2c). In addition, Notch1 + tumour cells appear more proliferative than non-marked tumour cells (Fig. 2d). We next assessed the cellular composition of Notch-derived clones and found that Notch1+ tumour cells can generate the repertoire of different cell types found within intestinal adenomas, as Notch1 lineages contain both proliferative (marked by Proliferating Cell Nuclear Antigen, PCNA in Fig. 2e) and differentiated tumour cells (marked by Lyz1, Muc2 and ChgA in Fig. 2e). The longevity of Notch1-expressing tumour cells, along with their multipotency, establishes that Notch1 is expressed *in vivo* in CSC.

**Figure 2.**
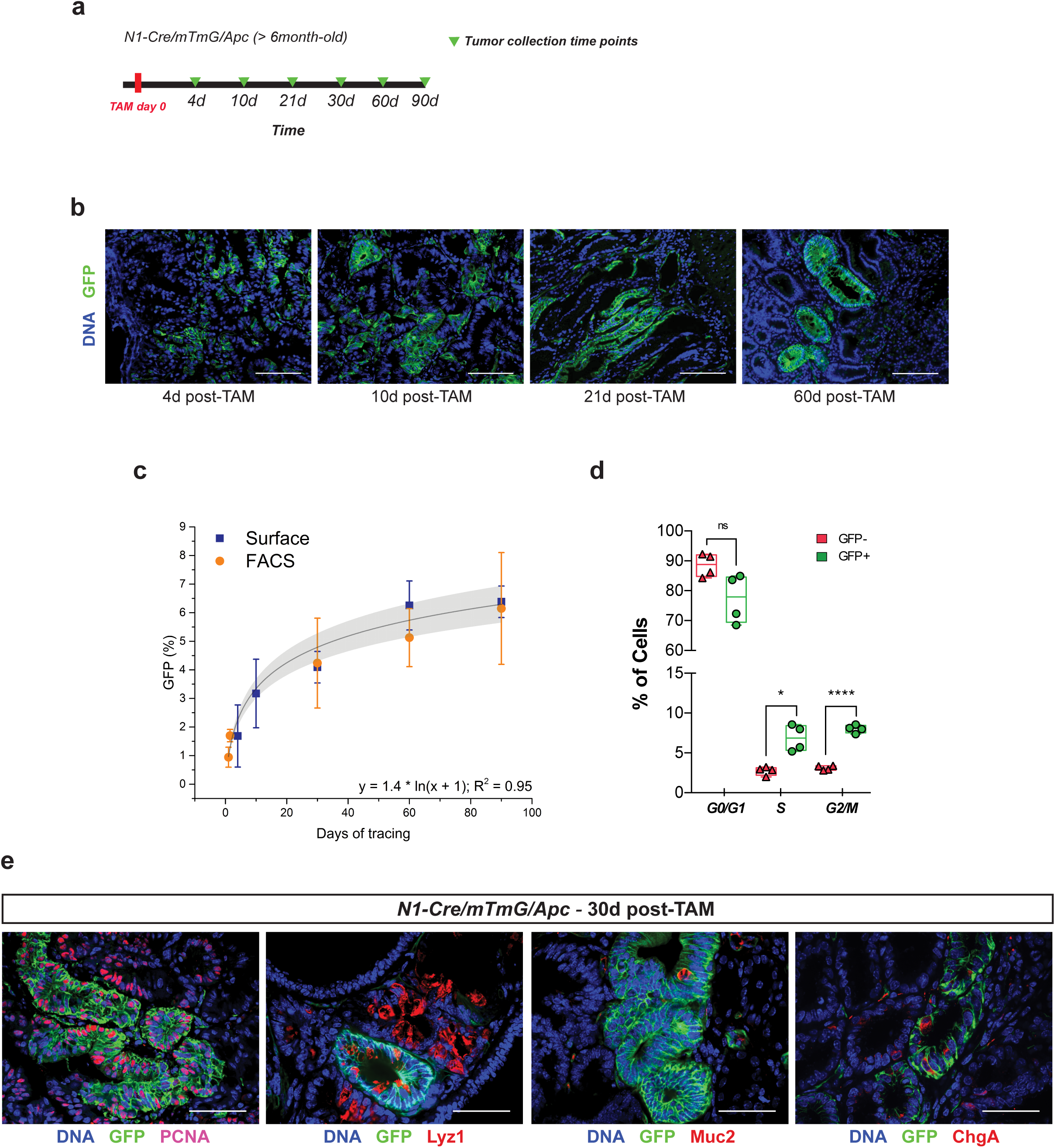
Notchl expression defines multipotent tumour cells with self-renewal capacity. **(A)** Experimental timeline used to perform lineage tracing analysis in N1-Cre/mTmG/Apc tumours *in vivo*. N1-Cre/mTmG/Apc tumour-bearing mice were induced at the age of about 6 months and then culled at 4, 10, 21, 30, 60 and 90 days. **(B)** Representative images of N1-Cre/mTmG/Apc tumour sections after 4, 10, 21 or 60 days chase showing the rapid generation of Notch1+ clones that grow over time. Notch1-derived lineages are labelled in green (GFP+) and DNA in blue by DAPI. **(C)** Clonal analysis of Notch1+ tumour cells using a dual methodology (FACS and fluorescence) demonstrates self-renewal capacity of Notch1+ tumour cells. Notch1-expressing GFP+ cells within TEC were quantified by FACS at the following tumour collection time points: 24h, 36h, 30d, 60d and 90d (orange dots; at least three biological replicates per time point). Quantification of the GFP+ tumour area within the total tumour surface (see also Supplementary Fig. 2c) was carried out at 4d, 10d, 30d, 60d and 90d post tamoxifen induction (blue squares). Both quantifications were best fitted by the y=1.4*ln(x+1) logarithmic function (R^2^ = 0.95). **(D)** FACS quantification of dividing (S and G2/M) and nondividing (G0/G1) TECs. Notch1+ tumour cells are represented by green dots and non-labelled cells by red dots. Boxes represent min to max values ± SDs of independent biological replicates (n = 4) ****= P ≤ 0.0005; **= P ≤ 0.005, *= P ≤ 0.05, using the Paired Ratio t-test. **(E)** Immunofluorescence analysis of tumour sections showing Notchl-derived lineages (GFP, in green) 30 days after induction using anti-PCNA, anti-lysozyme (Lyz1), anti-Mucin2 and antiChromogranin A, all in red. DNA is marked by DAPI in blue. Scale bars correspond to 100rfm in **(B)** and 50rfm in **(E)**.

### The transcriptional signature of Notch1+ tumour cells reveals a close correlation with the gene expression profile of normal ISCs

To molecularly characterize the tumour cells that express the Notch1+ receptor, we initially assessed the expression of selected genes in sorted cells by qRT-PCR. We confirmed that GFP+ cells expressed high levels of GFP, Notch1, Nrarp, Hes1 and Olfm4, the three-latter representing direct Notch targets^19-21^, (Fig. 3a), indicating that the Notch pathway is indeed active in Notch1+ tumour cells. Notch1+ tumour cells (GFP+) also showed enriched expression in reported markers of ISCs, such as Ascl2^22^, Hopx^2^, Musashi1 (Msi1)^23^ and Bmi1^7^ compared to GFP-cells within the same tumours (Fig. 3a). Next, we defined the genome-wide transcriptional signature of Notch1+ tumour cells and normal ISCs by performing Affymetrix analyses of FAC-sorted GFP+ and GFP-tumour and normal crypt cells derived from N1-Cre/mTmG/Apc and N1-Cre/mTmG mice, respectively. This analysis confirmed that both tumour cells and normal ISCs expressing the Notch1 receptor are undifferentiated, as they show downregulation of differentiation markers, such as Mucin2 (Muc2), the Regenerating Family Member 4 (Reg4^24^), Chromogranin A (Chga^25^), Doublecortin Like Kinase 1 (Dclk1^26^), Alpi and Membrane Alanyl Aminopeptidase (Anpep, an absorptive marker^27^), while they express high levels of reported ISCs markers, including Olfm4^28^, Lrig1^29,30^, Smoc2^31^, Hopx and Aldh1b1 (Aldehyde dehydrogenases 1B1^32^) (Fig. 3b).

**Figure 3.**
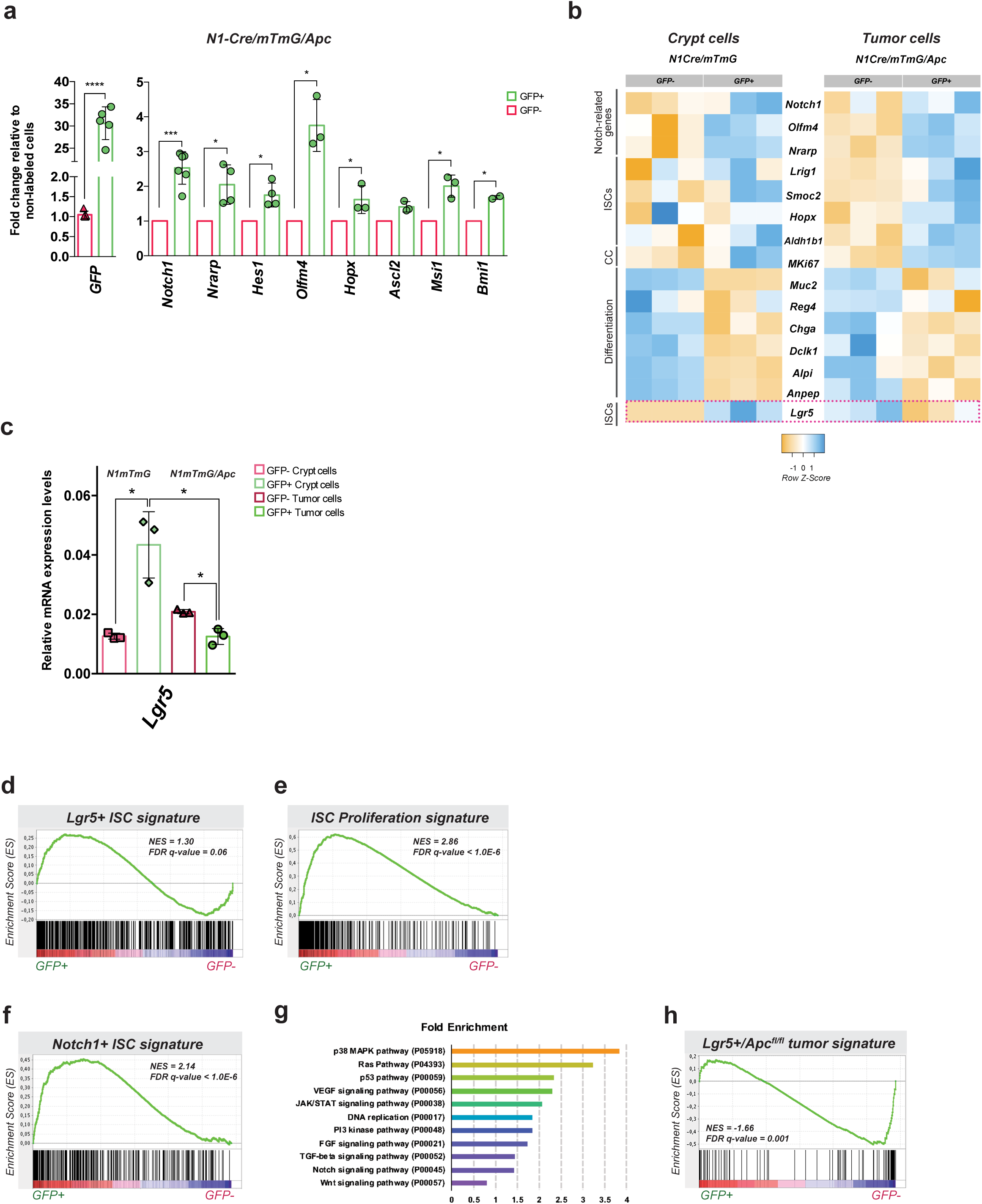
The transcriptional signature of Notch1+ tumour cells reveals a close correlation with the gene expression profile of normal ISCs. **(A)** RT-PCR for the indicated genes of sorted Notch1+ tumour cells (green bars) relative to non-recombined tumour cells (red bars). Data is expressed as the mean ± SDs of different biological replicates (each represented by dots) normalized by 18S housekeeping gene expression. ****= P ≤ 0.0005, **= P ≤ 0.005; *= P ≤ 0.05. The p-values were calculated using the Paired Ratio t-test. **(B)** Heatmaps showing expression patterns of selected genes extracted from genome-wide transcriptional datasets comparing Notch1+ crypt cells (GFP+ sorted cells from N1-Cre/mTmG mice) and Notch1+ tumour cells (GFP+ sorted cells N1-Cre/mTmG/Apc mice) relatively to non-labelled sorted crypt and tumour cells (GFP-). Expression levels of three independent biological replicates for each selected gene; Notchl, Notchl targets (Nrarp, Hes1 and Olfm4), ISCs markers (Lrigl, Smoc2, Hopx, Aldhlbl and Lgr5), proliferation antigen (MKi67) and differentiation markers (Muc2, Reg4, ChgA, Dclkl, Alpi and Anpep) are depicted by a scale of colour intensity measured by the Z-score (varying from ‐1 to 1). **(C)** RT-PCR showing the relative RNA expression levels of Lgr5 in Notch1+ crypt cells (GFP+ sorted cells from N1-Cre/mTmG mice, in light green) and Notch1+ tumour cells (GFP+ sorted cells from N1-Cre/mTmG/Apc mice, in dark green) relative to non-labelled cells (shown in light pink and dark pink). Data is shown as the average ± SDs of three independent biological replicates normalized by 18S housekeeping gene. *= P ≥ 0.05 using the Student’s t test (Welch correction). **(D-F)** Gene set enrichment analyses (GSEA) profiling positive correlations between Notchl-expressing tumour cells and normal Lgr5+ ISC signature^31^ **(D)**, ISC proliferative signature^33^ **(E)**, and Notch1+ ISC signature **(F)**. **(G)** PANTHER (Protein Analysis Through Evolutionary Relationships^42^) pathway analysis of signalling pathways altered in Notchl-expressing tumour cells compared to Notch1+ ISCs. The graph displays selected signalling pathways associated with a fold enrichment > 0.79. **(H)** Gene set enrichment analyses (GSEA) profiling positive correlation between non-labelled tumour cells and Lgr5+/Apc^fl/fl^ tumour cells signature^34^.

Unexpectedly, while in normal crypts the Notch1 and Lgr5 transcripts are enriched within the same cells, Notch1+ tumour cells showed a reduction in Lgr5 expression compared to nonlabeled cells (Fig. 3b-c).

The expression profile of Notch1+ tumour cells highly correlates, by Gene Set Enrichment Analysis (GSEA), with published transcriptomic signatures defining normal ISCs^31^ (Fig. 3d) and their proliferative features^33^ (Fig. 3e). Consistently, we also found a tight correlation between the cells that express Notch1 in normal crypts and in tumours (Fig. 3f). These results suggest that the transcriptional signature is not substantially changed between normal ISCs and CSCs, although we could detect a significant activation of proliferative and oncogenic pathways, such as MAPK, EGFR, FGF, VEGF and Wnt, in Notch1+ tumour cells compared to Notch1+ normal ISCs (Fig. 3g and full pathway analysis list in Supplementary Table 1). Collectively, these results indicate that Notch1-expressing tumour cells are transcriptionally related to normal ISCs. Based on the comparison between these two transcriptional signatures, it is tempting to speculate that Notch1+ CSCs originate from ISCs.

### Notch1 and Lgr5 lineages are hierarchically organized

Given our surprising results in terms of Lgr5 expression in Notch1+ cells, we then compared the signature of Notch1+ tumour cells to the transcriptome of Lgr5+ tumour cells^34^, and found an inverse correlation by GSEA, suggesting major differences between these two tumour cell populations (Fig. 3h). To explore the hierarchy between these two tumour cell populations, we used the mTmG double fluorescent line, allowing us to distinguish between initially labelled cells (that we called “mother cells”) and their derived progeny. Notch1-expressing cells are isolated 24h after tamoxifen administration; within such a short chase, Notch1+ cells have turned green (GFP+) but retain the expression of the Tomato protein, due to protein stability, so they are GFP+ and Tom+. By labelling Notch lineages in tumours during a 60 days chase and giving a second tamoxifen pulse 24h prior to tumour dissection (Fig. 4a), we can distinguish the recently labelled cells (“mother cells”, expressing both GFP and Tomato) from their progeny (GFP+ but Tomato-) and from non-labelled cells (GFP‐ and Tomato-) (Fig. 4b). The analysis by qRT-PCR of these three FACS sorted populations showed that Notch1-derived lineages express intermediate levels of both Lgr5 and differentiation markers, such as lysozyme, between non-labelled cells (in red) and Notch1-expressing tumour cells (in green) (Fig. 4c). These results are consistent with our clonal analysis, suggesting that multipotent Notch1+ tumour cells can give rise to heterogeneous lineages comprising distinct cell types, such as differentiated cells (Lyz1+), and raises the intriguing possibility that Notch1+ tumour cells might generate Lgr5+ tumour cells, thus contributing to tumour growth and propagation.

**Figure 4.**
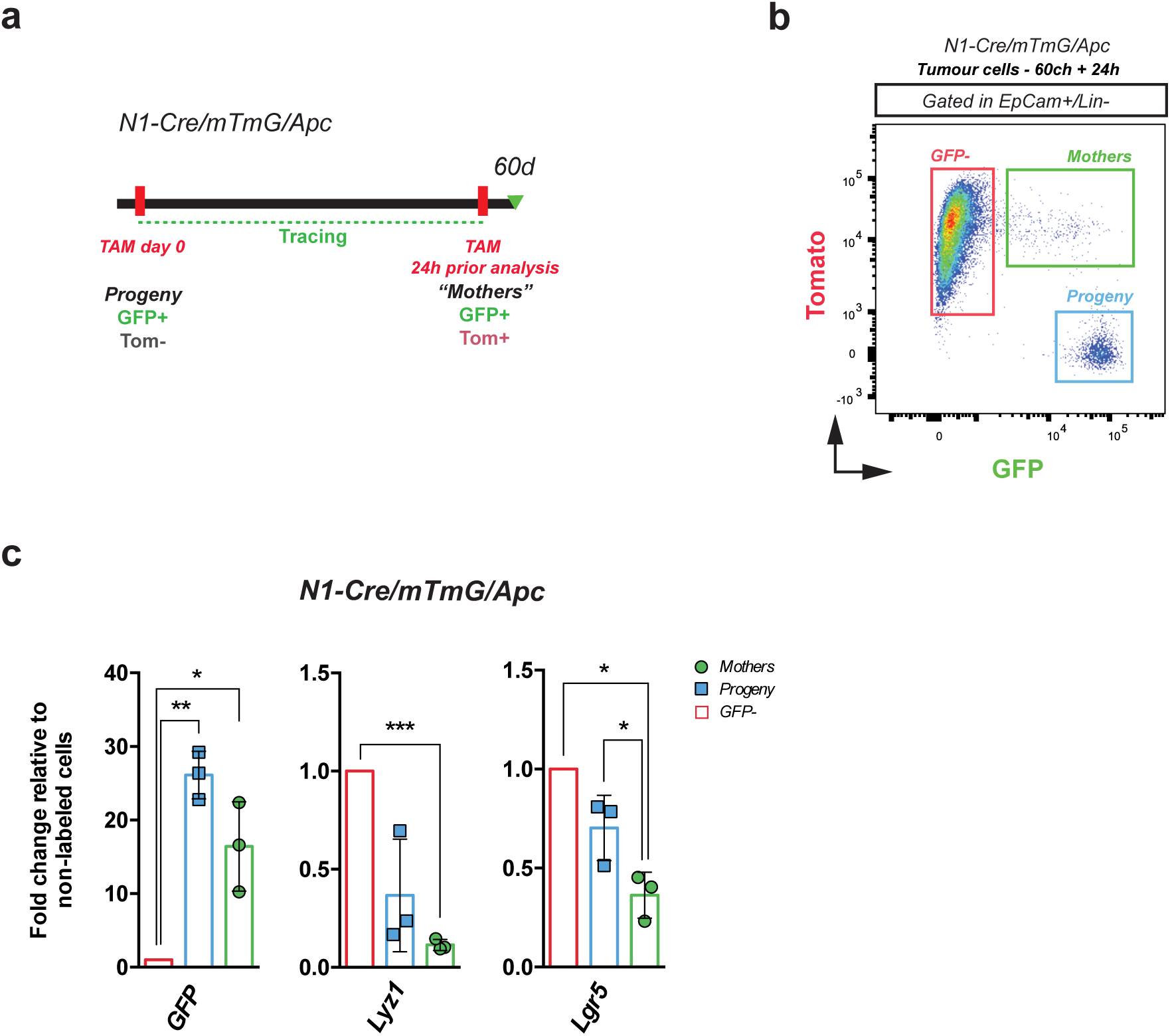
Notch1+ and Lgr5+ tumour cells are hierarchically organized. **(A)** Schematic representation of genetic lineage tracing strategy to mark Notchl-derived progeny and Notchl-expressing cells (“mothers”) within the same mouse. N1-Cre/mTmG/Apc tumour-bearing mice were induced with a single pulse of tamoxifen at day 0 and allowed to generate a clonal progeny for 60d (Progeny; GFP+/Tom-cells). 24h before culling, mice received a second injection of tamoxifen, in order to label Notchl-expressing tumour cells (GFP+/Tom+ cells). **(B)** Representative FACS dot plot of N1-Cre/mTmG/Apc tumour cells analysed upon a 60 days chase and a 24h re-pulse (according to the diagram in **A)**. Dissociated tumour cells were gated within TEC (Epcam+/Lin-) and Notch1+ tumour cells (“Mothers” highlighted in green; GFP+/Tom+), Notchl progeny (GFP+/Tom_ cells, in blue) and non-labelled tumour cells (GFP-/Tom-, in red) were analysed. **(C)** RT-PCR analysis showing mRNA expression levels of GFP, Lgr5 and Lysozymel (Lyz1) in Notchl-expressing tumour cells (Mothers represented by green bars) and Notchl-derived Progeny (Progeny displayed in blue), relative to GFP-(in red). mRNA expression was normalized to 18S. Data is expressed as the mean ± SDs of biological replicates (n = 3) normalized by 18S housekeeping gene expression. ***= P ≤ 0.005; **= P ≤ 0.005; *= P ≤ 0.05. The p-values were calculated using the Paired Ratio t-test.

### Clonal analysis of Notch1+ tumour cells in chemically induced colon tumours

Our clonal analysis in genetically modified mouse models indicated that the Notch1 receptor is expressed in a population of CSCs in intestinal adenomas. To further validate the clinical value of our study, we then examined if Notch1 expression could assist in identifying CSCs in chemically induced colon tumours, the most predominant location in colorectal cancer (CRC) patients. For these studies, we administered azoxymethane (AOM), followed by 2 cycles of exposure to the inflammatory agent dextran sodium sulphate (DSS)^35^ to N1-Cre/mTmG mice. Upon colon tumour formation, which we monitored by colonoscopy^36^, we induced GFP expression with a single dose of tamoxifen and analysed the tumours at different chase times (24h, 48h, 15 days and 2 months) (Fig. 5a). The presence of uniformly scattered GFP+ cells 24 hours post-tamoxifen in all colon tumours analysed demonstrated that Notch1 is also expressed in chemically-induced colon tumours (Fig. 5b). Consistently with our results in Apc mutant adenomas, Notch1+ colon tumour cells were also undifferentiated and proliferative, as they rapidly produced a clonal progeny, evident at 15 days after induction. A 2-month chase showed that Notch1-derived clones had considerably grown, suggesting that Notch1+ colon tumour cells self-renew, like their small intestinal counterparts (Fig. 5b).

**Figure 5.**
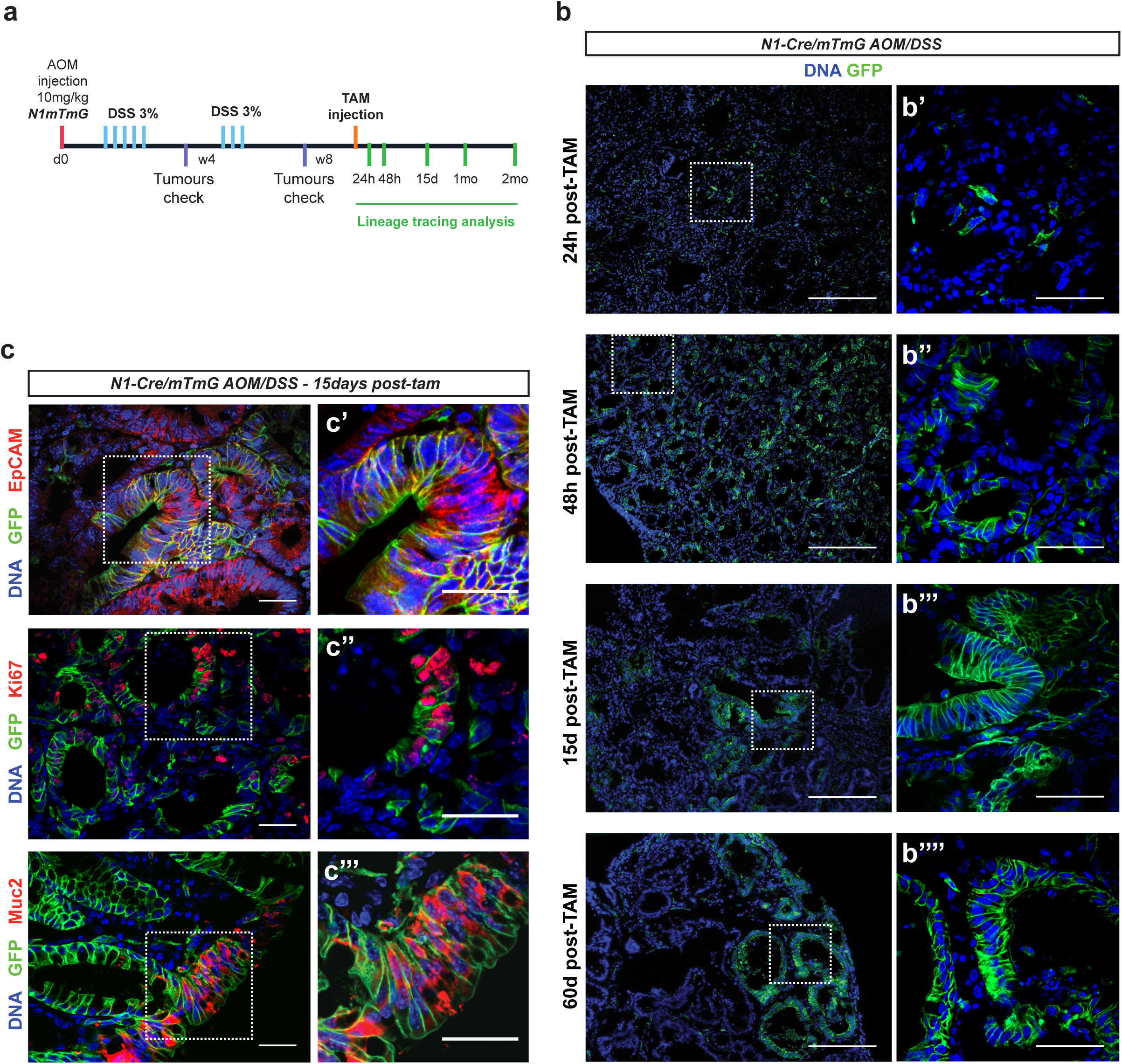
Clonal analysis of Notch1+ tumour cells in chemically induced colon tumours. **(A)** Experimental protocol and lineage tracing analysis in chemically induced colon tumours. N1-Cre/mTmG mice were injected with AOM at day 0 (d0) and received 2 cycles of 3% DSS (blue bars), spaced by 3 weeks of recovery. After 4w (4 weeks) and 8w (8 weeks) the development of colon tumours in control mice was monitored by colonoscopy. Lineage tracing analysis was started after 8 weeks and tumours were harvested at the indicated time-points (green bars). **(B)** Representative sections of colon tumours analysed at the indicated timepoints after tamoxifen administration. Notch1+ colon tumour cells and Notchl-derived clonal progeny present membrane GFP expression (in green). DNA is marked in blue with DAPI. **(C)** Immunostaining of tumour sections 15 days after Cre induction using anti-EpCAM, anti-Ki67 and anti-Mucin2 (all in red). Notch1+ cells are revealed by membrane GFP expression in green and nuclei in blue. Insets highlight areas of the tumour containing clones derived from Notchl-expressing colon tumour cells that co-localize with the used markers. Scale bars represent 200rfm in (B) and 50rfm in (B’-B’’’’), 30rfm in both (C) and respective high magnification panels (C’-C’’’).

To assess multipotency of the Notch1+ colon cancer cells, we analysed their clonal progeny (EpCAM+) and found that they expressed both proliferation (Ki67+) and differentiation markers (i.e. Mucin2 shown in Fig. 5c), demonstrating that Notch1-expressing cells in colon tumours have multilineage differentiation potential and present the same properties of Notch1+ intestinal adenoma cells. Notch1-expressing cells represent thus genuine CSCs in both genetic and carcinogenic tumour models. It remains uncertain however to what extent these cells contribute to the overall tumour development and growth.

## Discussion

Our study shows that Notch1+ expression labels a previously uncharacterized and distinct population of undifferentiated and self-renewing tumour cells, which clonally expand during tumour growth and generate heterogeneous lineages. Importantly, our findings suggest that Notch1+ tumour cells do not coincide with CSCs expressing the Lgr5 receptor, while they rather share a transcriptional signature with normal ISCs. In the normal intestine, two types of ISCs have been defined: rapidly dividing crypt-based columnar (CBCs) cells at the bottom of the crypt and “+4” ISCs (often called reserve stem cells), located at position +4 above the crypt base and considered as slower cycling cells^1,7^. We have previously shown that Notch1 expression marks both types of ISCs^4^. The fact that Notch1+ tumour cells present an expression profile very similar to the signature of Notch1+ ISCs (Fig. 3f), prompts us to speculate that Notch1+ tumour cells might derive from +4 ISCs, shown to express lower levels of Lgr5 than CBCs, and paramount for intestinal regeneration^3^.

It is noteworthy that our data show an inverse correlation between the transcriptional profile of Notch1+ tumour cells and the signature of Lgr5+ tumour cells published by Schepers and colleagues^34^. Such discrepancy might be due to the different models used in the two studies: Schepers et al. analysed Lgr5-expressing cells within intestinal tumours in which the homozygous deletion of the *Apc* gene had been specifically targeted to Lgr5+ cells^34^. This approach greatly differs from our study, where we traced Notch1+ cells in spontaneously arising tumours that lost the wt *Apc* allele by loss of heterozygosis (LOH). In this model, we found that Notch1+ tumour cells strikingly resemble their normal counterparts, inferring that they might represent normal stem cells that were engulfed during tumour growth and thrive in the tumour environment thanks to paracrine mitogenic signals from surrounding cells (Fig. 6). Our work suggests the possibility that normal ISCs might contribute to tumour expansion and heterogeneity. We also show that, independently of their origin, Notch1+ tumour cells share many features with normal ISCs, stressing the complications in developing new CSC-targeted drugs. The transcriptomic signatures presented in our study provide the means to identify novel biomarkers within the Notch1 + CSC population that might allow the distinction between normal and tumour stem cells in future drug screens.

**Figure 6.**
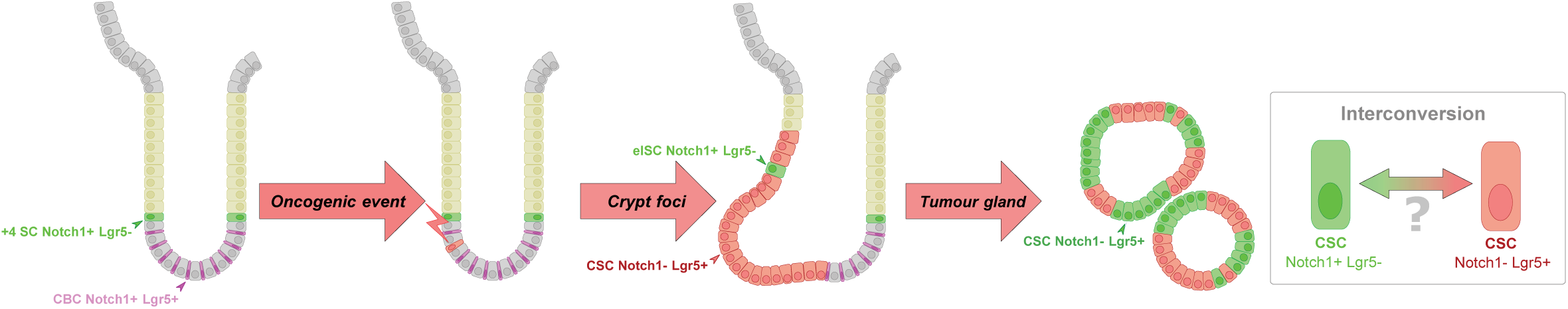
Proposed model for a cellular hierarchy within spontaneous tumours. In homeostatic conditions, Notch1 is expressed in both types of ISCs, the +4 SCs (in green), and the CBC ISCs, enriched in Lgr5 expression (in purple). According to the bottom up model for tumourigenesis, an oncogenic hit, such as loss of the tumour suppressor *Apc*, should occur in an ISC to initiate crypt hyperplasia (crypt foci, in red). In such a scenario, Notch1+ normal ISCs become surrounded by mutant tumour-initiating cells (red cells), and thus get engulfed in the tumour. As mutant cells expand, their tumour microenvironment provides signals that potentiate the growth of these “embedded ISCs” (eISCs, in green), which express lower levels of Lgr5 relatively to normal ISCs and to the tumour bulk (see Fig. 3c). As the adenoma continues to develop over time, eISCs remain proliferative and multipotent within tumour glands, actively producing differentiated cells and generating new Notch1+ CSCs, but also Lgr5+ CSCs (red cells) that display lower levels of Notch1, as we speculate (see Fig. 4c). The existence of two types of CSCs, one predominantly expressing Notch1 and the other Lgr5, might contribute to intra-tumoural heterogeneity, which drives tumour progression, by escaping to targeted therapies. Whether Lgr5+ CSCs are reciprocally able to produce/interconvert into Notch1+ CSCs remains to be established.

Of relevance, our results define a cellular hierarchy within tumours, with Notch1+ CSCs possibly giving rise to Lgr5+ tumour cells. Moreover, given the well-established plasticity and interconversion of different cell types in the normal intestine^2,3^, it is possible that Lgr5+ tumour cells could also generate Notch1+ cells, though only Lgr5 clonal analysis in the same mouse model could unambiguously establish if this is the case. Notably, a recent study demonstrated that intestinal tumours comprise distinct pools of functionally different CSCs^37^, that reciprocally and dynamically interconvert, corroborating our results on the existence of two distinct CSCs populations, marked by Notch1 or Lgr5 (Fig. 6). Moreover, CRISPR-mediated tracing of Lgr5+ CSCs in human CRC organoids demonstrated the re-emergence of newly generated Lgr5+ CSCs upon their targeted ablation, revealing widespread plasticity of human CRC cells^38^. Our work proposes the existence of distinct types of CSCs that can dynamically interconvert, a concept that anticipates an extra layer of complexity in developing drugs targeting specific tumour cell populations. Whereas selective ablation of Lgr5+ CSCs does not lead tumour regression, but only temporarily reduces tumour growth^37,38^, it remains to be established whether deletion of Notch1+ CSCs would be beneficial to block or retard tumour progression.

Further functional studies will reveal the differences between the distinct CSCs populations revealed in this study, in particular regarding their respective clonogenic and tumourigenic potential. It will also be crucial to test their resistance to commonly used chemotherapeutic drugs, to possibly stratify patients based on their relative abundance.

## Methods

### Mice

The N1-CreERT2^SAT 4^ and Apc^+/1638 14^ mouse lines have been crossed to the dual fluorescent reporter Rosa26mT/mG strain^13^. GFP reporter expression was induced in N1-Cre/mTmG/Apc 6-8 month-old mice or in N1-Cre/mTmG adult mice by intraperitoneal injection of 0.1mg/g of mouse body weight of tamoxifen free base (Euromedex). Mice were culled either 24h post-induction (Notch1-expressing cells) or at different time points ranging from 36h up to 3 months after induction. For each time point, at least three mice displaying tumours were analysed. No GFP fluorescence was observed in non-induced mice.

### Ethics Statement

All studies and procedures involving animals were in strict accordance with the European and National Regulation for the Protection of Vertebrate Animals used for Experimental and other Scientific Purposes. The project was approved by the French Ministry of Research with the reference #04240.03 and was fully performed in the Animal Facility of Institut Curie (facility license #C75–05–18). We comply also with internationally established principles of replacement, reduction, and refinement in accordance with the Guide for the Care and Use of Laboratory Animals (NRC 2011). Husbandry, supply of animals, as well as maintenance and care of the animals in Exempt Of Pathogen Species (EOPS) environments before and during experiments fully satisfied the animal’s needs and welfare. Suffering of the animals has been kept to a minimum.

### Tissue dissociation into single cells

Freshly dissected tumours were dissected in DMEM/F12 (2% PS), following incubation in 5mM EGTA-PBS at 4°C, shaking gently, to remove potential contaminant cells from the normal tissue. Tumours were then minced in small pieces with razor blades and incubated in diluted TrypLE Express in PBS, shaking (180rpm) for 45 minutes at 37°C. Trypsin was inactivated with 10% of cold FBS. The cell suspension obtained was then filtered through a 40μm cell strainer and cells were counted upon centrifugation for 5 min at 450g, following resuspension in the appropriated medium. Small intestines were harvested and flushed with cold 1X PBS, following longitudinal opening and cut into small pieces of approximately 2mm × 2mm. Intestinal fragments were subsequently incubated with 2mM EDTA in HBSS for 30 minutes at 4°C. Crypts were obtained by serial fractioning, following TrypLE Express (ThermoFisher Scientific) incubation for 5 minutes at 37°C to obtain single cells. TrypLE

Express was inactivated with 10% cold FBS (Sigma-Aldrich). To confirm crypt separation, 20 μl of supernatant sample was checked under the microscope. The cell suspension obtained was then filtered through a 40rfm cell strainer and cells were counted upon centrifugation for 5 min at 450g, following suspension in Flow buffer.

### Immunofluorescence

Freshly dissected intestines and tumours were washed in 1X PBS and fixed at room temperature with 4% PFA under agitation for 2h. The samples were then impregnated with 30% Sucrose (for at least 24h at 4°C) or dehydrated in 70% Ethanol and embedded in OCT (VWR) or in paraffin, respectively. Samples were sectioned either in a cryostat or microtome at 5μm thickness. For immunofluorescence staining, frozen sections were incubated with 0,3% Triton-blocking buffer (5% FBS and 2% BSA). Paraffin-embedded sections were rehydrated through a gradient of Ethanol. Subsequently, antigen retrieval was achieved by boiling in 10mM citrate buffer (20 minutes) for all antibodies. The following primary antibodies were used: chicken anti-GFP (1:800, ab13970), rabbit anti-Lysozyme (1:500, Dako A009902), rabbit anti-Chromogranin A (1:200, Immunostar 20086), rabbit anti-Mucin2 (1:100/200, Clone PH497), mouse anti-PCNA (1:1000, a29), rabbit anti-Ki67 (1:200, ab15580), rabbit anti-EpCAM (Abcam, ab32392). Secondary antibodies were incubated in PBS for one to two hours at room temperature. The following secondary antibodies were used: anti-chicken AlexaFluor488 (1:500, Invitrogen A-11039), anti-rabbit AlexaFluor633 (1:500, Invitrogen A-21071), anti-mouse AlexaFluor633 (1:500, Invitrogen A-21202), antimouse Cy3 (1:500, Jackson laboratories 92557), anti-rabbit Cy3 (1:500, Jackson laboratories 91144). Identification of secretory cells on frozen sections was performed using Ulex Europeus Agglutinin I (UEA) directly coupled to Cy3 (1:50, Sigma-Aldrich). DNA was stained with DAPI.

### Quantification of clonal expansion by immunofluorescence

Whole tumour sections of 4-5μm derived from mice culled at different time points after tamoxifen injection were imaged for GFP (recombined tumour cells) and Tomato endogenous fluorescent proteins. To quantify the GFP surface area over the total tumour area, we used an unbiased approach taking advantage of the ImageJ Software threshold tool. By adjusting the threshold of each individual channel (GFP and Tomato), we extracted a binary image that contained exclusively black and white pixels and quantified these using a macro developed for ImageJ software. The sum of the Tomato pixels reflected the total area of the tumour (∑ pixels TOM; hence referred as A) and the sum of the GFP pixels indicated the recombined areas (∑ pixels GFP; called B). Since we have used homozygous mice carrying two copies of the Rosa26mTmG allele, resulting in GFP+/Tomato+ cells, GFP+ areas were subtracted from total (Tomato+) areas. The final area (A - B) was set to 100% and GFP area was calculated by the equation: GFP (%) = B*100% / (A - B).

### Fitting curve

The concatenated fit of the GFP cells expansion data from FACS and the surface-based methods was obtained using OriginLab software (OriginLab Inc., MA, USA). To account for the initial rapid increase and later slower growth of the GFP population, a logarithmic function has been chosen, and the quality of the fitting was judged by visual inspection and analysis of fit residuals and proved more appropriate than using an exponential decay function.

### Cell cycle analysis by FACS

After dissociation, cells were fixed in 2% PFA for 20 minutes at 4°C. Cells were washed twice in 1X PBS upon centrifugation at 400g, 4 minutes at 4°C, and staining proceeded. Cells were then incubated in pre-warmed 1X PBS Hoechst 33342 (1:100, stock at 10mg/ml) solution for 20 minutes at 37°C. FlowJo software was used for data analysis.

### Microscopy and image acquisition

For image acquisition, we used an upright confocal spinning disk (Roper/Zeiss), equipped with a CoolSnap HQ2 camera and 405nm, 491, 561 and 642nm lasers. Images were captured using Metamorph. Image processing was performed using ImageJ software.

### Flow cytometry

Dissociated cells were incubated in Flow buffer (DMEM/F12, 5mM EDTA, 1% BSA, 1% FBS and 10U/ml DNAse) during 25 minutes at 4°C with the following antibodies: EpCAM-PE/Cy7 (PE; R-Phycoerythrin, Cy7; Cyanine, 1:100, Biolegend clone G8.8), CD45-APC (APC; Allophycocyanin, 1:100, Biolegend clone 30-F11), CD31-APC (1:100, Biolegend clone MEC13.3), Ter-119-APC (1:100, Biolegend clone TER-119). To exclude non-viable cells, DAPI (1:1000, Sigma-Aldrich) was added. Cells were then washed and filtered directly into FACS tubes (40 μm strainer). Analysis was carried out on a FACS-LSRII and sorting on a FACS-Aria III (Becton Dickinson). For cell sorting, RTL lysis buffer supplemented with Beta-mercaptoethanol was used for RNA extraction (Qiagen). FlowJo software was used for data analysis.

### RNA extraction and qRT-PCR

RNA extraction was performed using Qiagen kit according to the manufacturer’s instructions. Reverse transcription was performed using the SuperScript III First-Strand Synthesis System (ThermoFisher Scientific), according to manufacturer’s instructions. Random hexamer primers were used for reverse transcription. Real-time PCR quantification of gene expression was systematically performed in triplicate using SYBR Green I Master (Roche) on a ViiA 7 RT-PCR System (ThermoFisher Scientific). The efficiency of the primers used for real-time quantification (listed in Supplementary Table 2) was evaluated relatively to the slope obtained by the quantification of a standard curve, and the presence of a single amplicon at the expected size was checked on an 2% agarose gel. Results were normalized to the expression of 18S, GAPDH and β-actin housekeeping genes and the relative expression was obtained with the ‐ΔΔCt method.

### AOM/DSS colon carcinogenesis experimental protocol

To induce colon carcinogenesis, we adjusted the protocol from Tanaka and colleagues^35^ and a set of N1/mTmG mice ranging from 5 to 7 months of age were administered a single intraperitoneal injection of Azoxymethane (AOM, Sigma #A5486) followed by Dextran Sulfate Sodium (DSS, MP Biomedicals #160110) treatment (3% in drinking water) the day after the AOM injection for 5 consecutive days. General health status and mouse body weight were monitored daily during and after treatment. To verify the presence of colon tumours, 2 mice were checked 1 month after the first cycle of DSS treatment, but no tumours were detected (only signs of inflammation). We administered another cycle of DSS (3% in drinking water) for 3 days and tumour formation was monitored by colonoscopy using a Karl-Storz endoscopic system (as previously described^36^).

### Affymetrix analyses and sample quality control

Transcriptome profiles were obtained using Affymetrix Mouse Gene ST 2.1 arrays. For each condition, three biological replicates were performed, by pooling intestinal adenomas and crypt fractions from more than 3 mice per replicate. cDNAs were synthesized from total RNA and hybridized by the Institut Curie Genomics Platform. Data normalization and analysis were performed using the R software (version 3.3.2). Raw data have been first normalized using the RMA method (R package affy 1.5.2, ^39^), and annotated using the BrainArray *Mus musculus* mogene21st annotation library (19.0.2). Non-annotated or lowly expressed genes (<3.5) were discarded from the analysis.

Differential analysis were performed using the Limma package (3.24.3)^40^ and the following linear model: Y_ij_ = mu_i_ + T_t_ + G_g_ + (T*G)_tg_ + E_ijtg_ where Y_ij_ is the expression of gene *i* in sample *j*, T is the effect of sample type ^41^ and G is the effect of GFP status. Results of the differential analysis have been corrected from multiple testing using the Benjamini-Hochberg procedure. Differentially expressed genes in the comparison of Normal GFP+ (Notch1+ ISCs) vs Tumour GFP+ (Notch1+ tumour cells), and Tumour GFP+ vs Tumour GFP-(non-labelled tumour cells), have been defined using an adjusted p-value of 5% and a minimum log fold change of 1.

We then performed GSEA analysis (with -nperm 1000 -permute gene_set -collapse false parameters) using the same expression matrix (v2.2.2) comparing the Tumour GFP+ and GFP-samples ^41^. Transcriptional signatures used for the analysis were extracted from the literatures ^31,33,34^. Normal GFP+ vs – signature have been defined using an adjusted p-value threshold to 25% and a minimum log fold change of 0.5.

PANTHER pathway analysis was used to determine enriched pathways in Notch1-expressing tumour cells vs Notch1+ ISCs ^42^.

The integrity of RNA samples used for transcriptomic analysis and qPCR analyses presented in this work was evaluated with a Bioanalyzer using the RNA 600 Pico lab chip (Agilent) according to manufacturer’s instructions. The amount and integrity of RNA samples selected for Affymetrix microarray was assessed with a Bioanalyzer and 2200 Tapestation system (Agilent) and the RNA integrity number (RIN) was confirmed to be higher than 8 for all the six samples. All transcriptomic experiments were performed using three biological replicates, consisting of cells pooled from at least 4 mice per replicate.

## Data-availability statement

Affymetrix data sets have been deposited in the Gene Expression Omnibus (GEO) database (pending accession code). All additional relevant data are available in the manuscript and its supplementary information files, or from the corresponding author upon request.

## Statistical analysis

Statistical analysis was performed using GraphPad Prism 7 (GraphPad Software, San Diego, CA, USA). P-values, calculated by Paired Ratio and Welch’s t-tests correspond to *= *p* ≤ 0.05, **= *p* ≤ 0.005; ***= *p* ≤ 0.0005; ****= *p* ≤ 0.0005. n ≥ 3 independent experiments were used for statistical analysis, unless otherwise indicated.

## Author contributions

S.F. and L.M. conceptualized and designed the experiments. L.M. performed experiments, analysed and interpreted the data. G.J. interpreted data and performed data visualization. W.J.N conducted formal data analysis. N.M. and N.S. analysed the transcriptomic data. M.H. provided technical support and contributed with unpublished data. L.M., G.J. and W.J.N prepared the figures. L.M. and S.F. wrote the manuscript. S.F. supervised the experiments.

## Acknowledgements

We are very grateful to Prof. Spyros Artavanis-Tsakonas for generously sharing the N1-Cre^ERT2^ mice. Likewise, we wish to acknowledge Prof. Riccardo Fodde and Prof. Shahragim

Tajbakhsh for kindly providing the Apc1638 mice and the mTmG reporter line, respectively. We would like to acknowledge the Cell and Tissue Imaging Platform PICT-IBiSA (member of France–Bioimaging) of the Genetics and Developmental Biology Department (UMR3215/U934) of Institut Curie for help with light microscopy; the Flow Cytometry and Cell Sorting Platform at Institute Curie for their expertise, in particular Zosia Maciorowski; the *In vivo* Experimental Facility, mainly Sonia Jannet, for help in the maintenance and care of our mouse colony; the Experimental Pathology facility at Curie Hospital for paraffin sample preparation and the Genomics Platform at Institut Curie for the transcriptomic analysis, in particular Audrey Rapinat and David Gentien. We particularly thank Audrey Michaud for technical help, Raphaël Margueron and Joan Barau for constructive discussions and experimental advice.

---

**Supplementary Figure 1.**
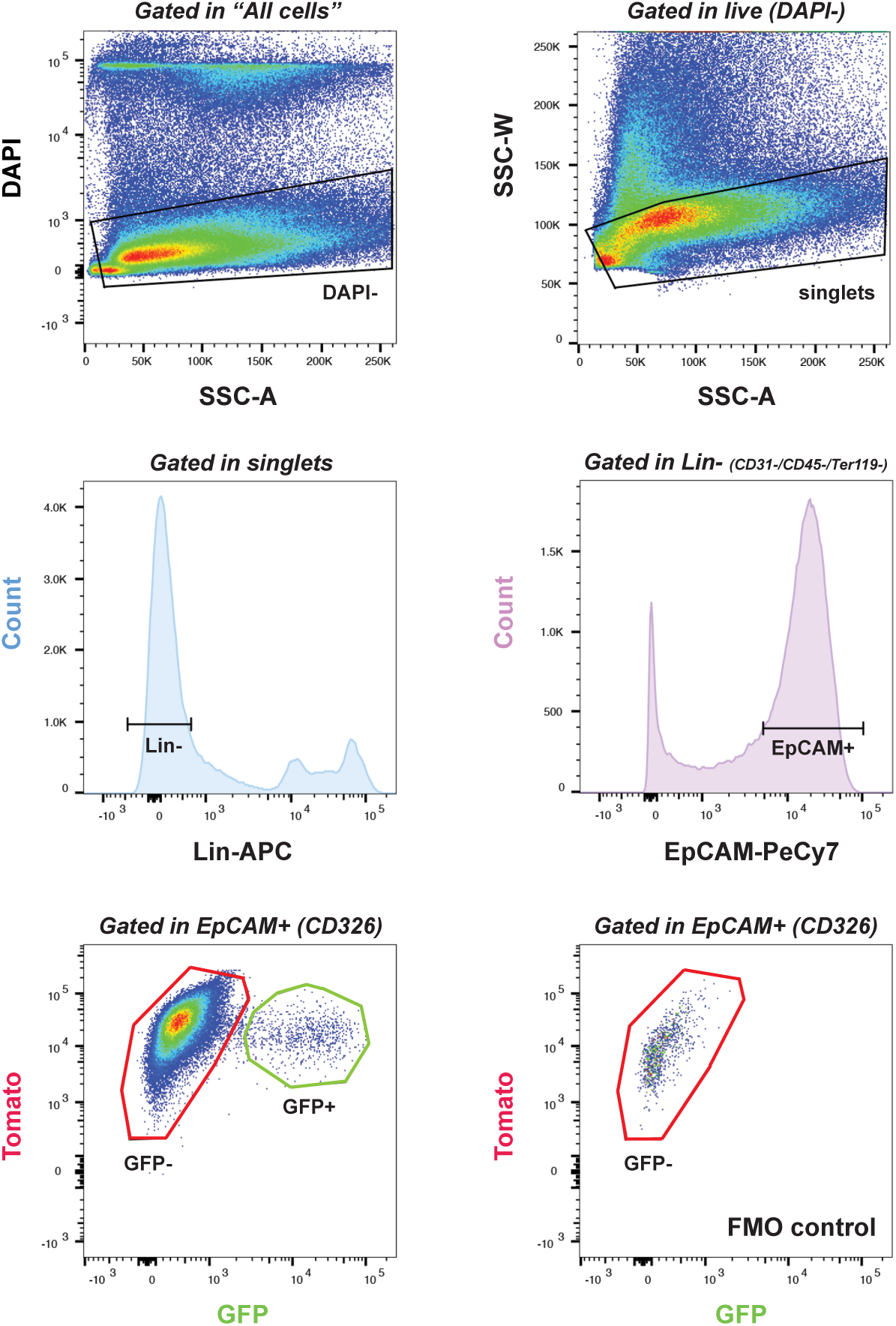
Gate strategies for FACS analysis. Representative dot plots of the FACS gating strategies used in this study. Live cells were gated in Dapi-/SSC-A. To remove cell aggregates, singlets were gated in SSC-W vs SSC-A. The sample was further analysed by selecting the Lin-population (Lin markers; CD31, CD45, Ter119), following for their uptake of the EpCAM (CD326) marker, resulting in the TEC (Tumour Epithelial Cells) gate. Notch1-expressing cells were gated using the GFP/Tomato channels. FMO (fluorescence minus one) control was used to draw GFP+ gates.

**Supplementary Figure 2.**
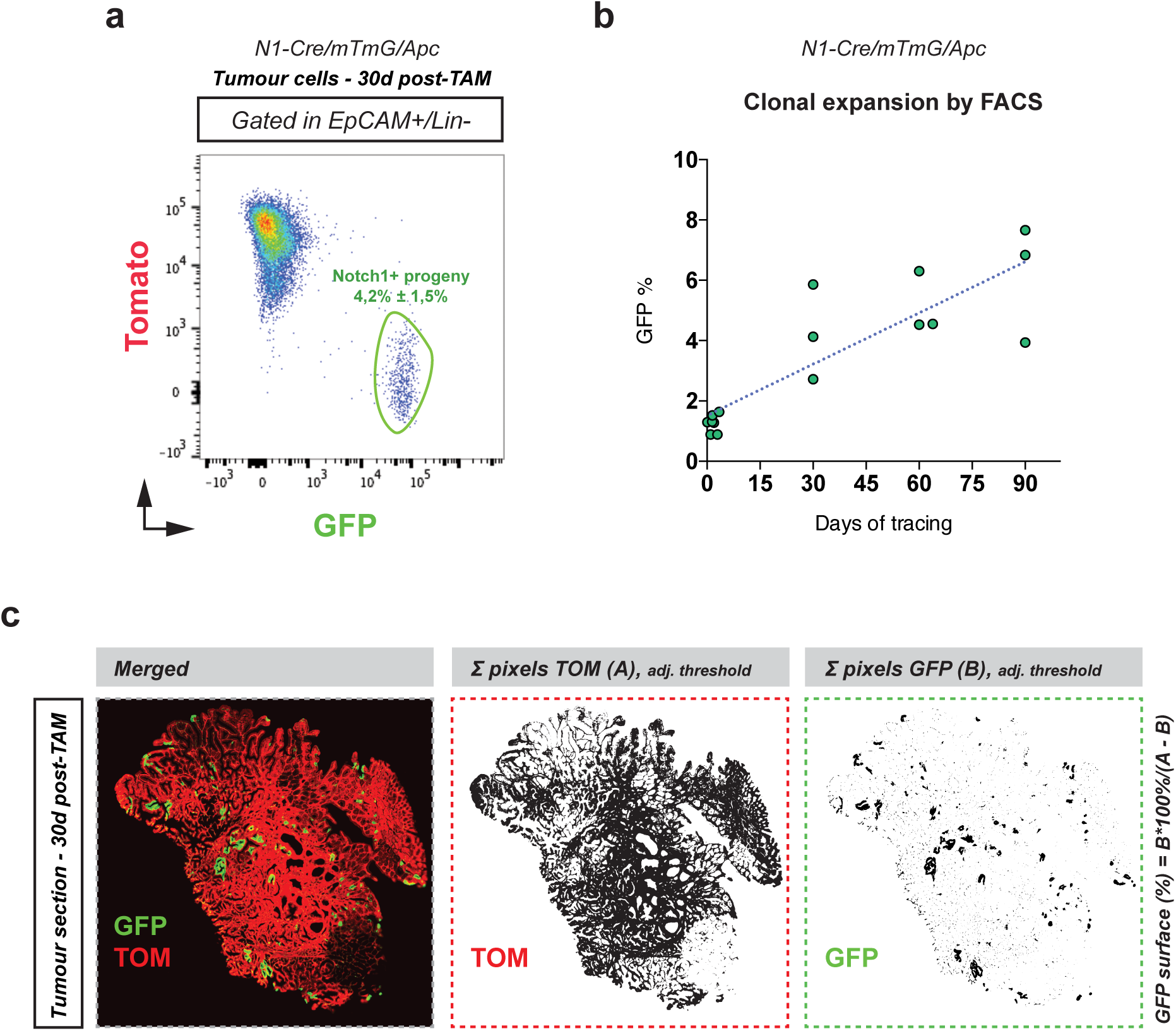
Quantification of clonal expansion by FACS analysis and immunofluorescence. **(A)** Representative FACS dot plot of N1-Cre/mTmG/Apc tumour cells gated as TEC and analysed upon a 30 days chase. The Notch1-derived progeny (GFP+/Tom-) no longer presents Tomato fluorescence. **(B)** Non-fitted FACS quantification of the clonal expansion of Notch1-expressing tumour cells. Each dot represents an independent biological replicate (n ≥ 3 per time point). **(C)** Schematic representation of the quantified GFP+ area in intestinal tumours. Images show a section of a tumour exhibiting endogenous Tomato (TOM) fluorescence (in red) and clones derived from Notch1-expressing tumour cells displaying GFP fluorescence (in green). Split channels of binary composites for both TOM (A representing the number of pixels present within the TOM channel) and GFP (B; representing the number of pixels present within the GFP channel) were used to quantify the GFP+ area.

